# Mechanistic insights into the interactions between cancer drivers and the tumour immune microenvironment

**DOI:** 10.1101/2023.01.24.525325

**Authors:** Hrvoje Misetic, Mohamed Reda Keddar, Jean-Pierre Jeannon, Francesca D. Ciccarelli

## Abstract

The crosstalk between cancer and the tumour immune microenvironment (TIME) has attracted significant interest because of its impact on cancer evolution and response to treatment. Despite this, cancer-specific tumour-TIME interactions and their mechanisms of action are still poorly understood. Here we identified the interactions between cancer-specific genetic drivers and anti- or pro-tumour TIME features in individual samples of 32 cancer types. The resulting 477 TIME drivers are multifunctional genes whose alterations are selected early in cancer evolution and recur across and within cancer types. Moreover, the anti-tumour TIME driver burden is predictive of overall response to immunotherapy. Focusing on head and neck squamous cancer (HNSC), we rebuilt the functional networks linking specific TIME driver alterations to the TIME state. We showed that TIME driver alterations predict the immune profiles of HNSC molecular subtypes, and that deregulation of keratinization, apoptosis and interferon signalling underpin specific driver-TIME interactions. Overall, our study provides a comprehensive resource of TIME drivers giving mechanistic insights into their immune-regulatory role.

## INTRODUCTION

Cancer evolves within a stromal microenvironment with whom it engages in a dynamic crosstalk whereby genetic alterations in the cancer cells modulate the microenvironment and, in turn, the microenvironment sculpt the cancer genome^1–3^. Besides shaping cancer evolution, tumour-stroma interactions, especially with the tumour immune microenvironment (TIME), impact on overall prognosis and response to treatments including immunotherapy^4,5^. Unravelling cancer-TIME interactions is therefore crucial to fully understand cancer biology.

Tumour-TIME interactions often involve genes that drive cancer evolution (cancer drivers). For example, loss-of-function (LoF) alterations in *TP53* reduce the antitumour infiltration of natural killer cells^6^ while gain-of-function (GoF) alterations in *KRAS* promote pro-tumour infiltration of myeloid-derived suppressor cells^7^. Moreover, deregulations of the WNT and PI3K-AKT cancer pathways result in CD8^+^ T cell exclusion^8^ and regulatory T cells increase^9^, respectively.

Recently, systematic genetic screens have expanded the repertoire of genes that can modulate cancer immune response. A preferential loss of tumour suppressors has been observed in mice with a functional immune system where they likely promote immune escape^10^. Moreover, genome-wide CRISPR screens in co-cultures of cancer and cytotoxic T cells have identified genes losses conferring resistance to T cell-mediated killing^11^. Although these screens enable identification of TIME-interacting genes beyond cancer drivers, they rely on cell or animal models rather than human samples and have so far assessed the TIME role of LoF alterations neglecting that of GoF alterations.

Large cancer genomic and transcriptomic datasets allow to compute tumour-TIME associations in pan-cancer cohorts and are unbiased towards the alteration type. These studies have so far reported a prevalence of *PDL1* amplifications in immune-hot tumours^12,13^ as opposed to in *APC, KRAS, IDH1* or *FGFR3* mutations in immune-cold tumours^12–15^. They have so far focused mostly on anti-tumour immunity relying on the same list of drivers applied to the whole pan-cancer cohort. The absence of any further filtering on the actual cancer-specificity of the driver activity likely led to falsepositive associations. Moreover, very little is still known on the molecular mechanisms of the tumour-TIME associations.

To overcome these limitations, here we have computed the interactions between manually curated and cancer specific lists of drivers and five anti- and pro-tumour immune features of 6921 samples in 32 solid cancers. We have then investigated the properties of the resulting genes and their potential to predict response to immunotherapy. Taking head and neck squamous cancer (HNSC) as an example, we have rebuilt the tumour-TIME functional networks in the three HNSC subtypes. This enabled us to unravel the mechanisms linking driver alterations to the TIME modifications at the individual sample level.

## RESULTS

### TIME drivers are multifunctional genes commonly altered across cancers

To identify the cancer genes interacting with the TIME (TIME drivers), we derived a reliable set of genes specifically contributing to the evolution of each of the 32 TCGA cancer types (**Fig. S1A**). We started from a pan-cancer collection of experimentally validated (canonical) and computationally predicted (candidate) drivers^16^ and assigned drivers to each cancer type according to an expert annotation of the literature. We then retained only drivers with damaging alterations in in 7,730 TCGA samples with matched genomic and transcriptomic data. We considered LoF alterations in tumour suppressors, GoF alterations in oncogenes, and both types of alterations in drivers with unclassified roles. Finally, we removed rarely damaged drivers for which no reliable associations could be computed. The final list included 254 canonical and 977 candidate drivers with damaging alterations in 6,921 samples across the 32 cancer types (**Fig. 1A**, **Table S1**).

**Fig. 1.**
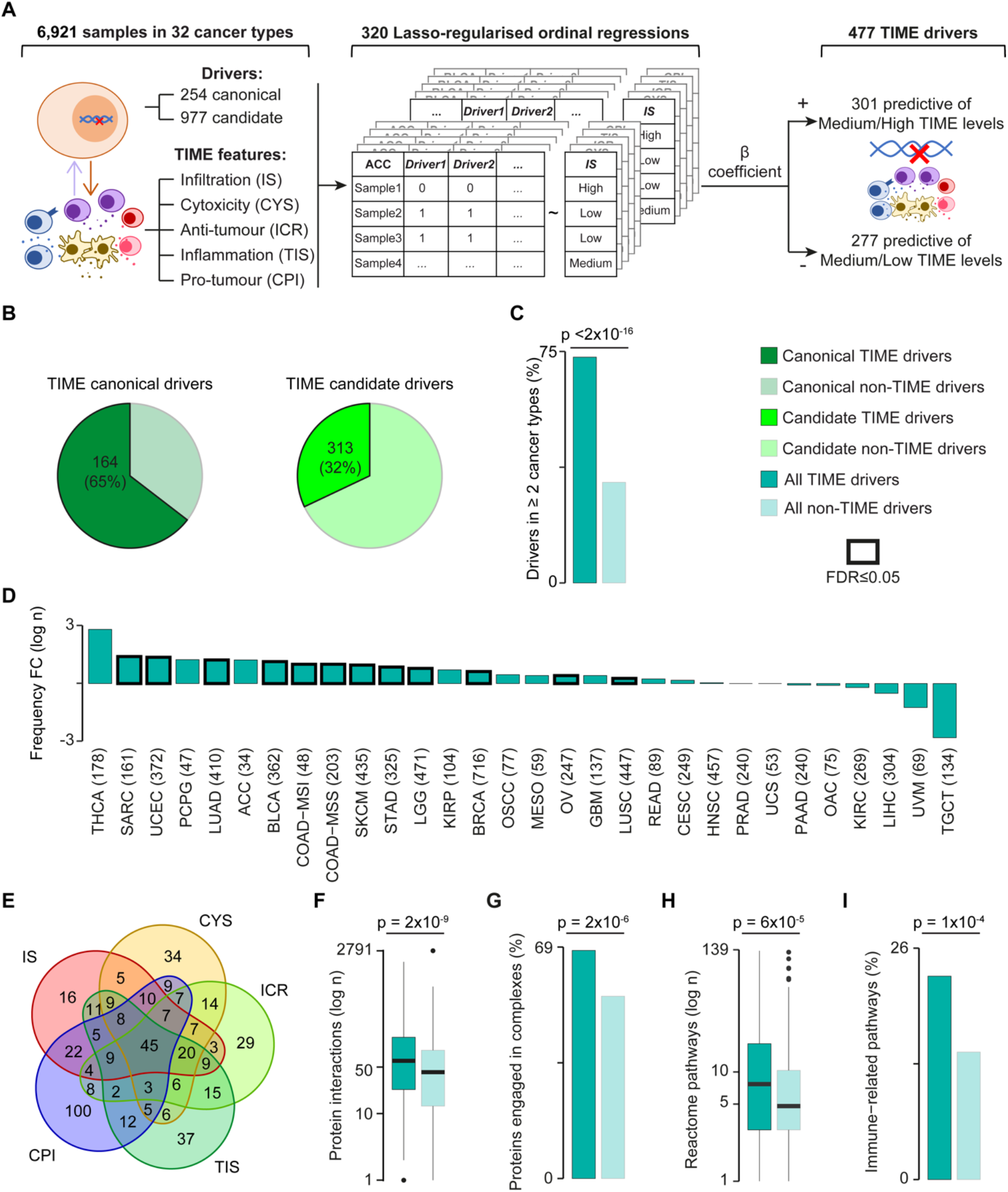
Identification and properties of TIME drivers. **A.** Approach for TIME driver prediction. Associations between damaging alterations in cancer-specific drivers and TIME features were assessed in 32 cancer types using Lasso-regularised ordinal regression. Regressions were computed separately for canonical and candidate drivers and TIME features in each cancer type. *β* >0 and *β* <0 indicated that altered cancer drivers were predictive of medium/high or medium/low TIME levels, respectively. **B.** Proportions of TIME canonical and candidate drivers over all drivers. **C.** Proportions of TIME and non-TIME drivers in ≥ 2 cancer types. **D**. Fold change (FC) of median frequency alterations of TIME and non-TIME drivers in each cancer type. The number of samples in each cancer type is shown in brackets. **E**. Venn diagram of TIME drivers predictive of the five TIME features. **F.** Distributions of protein-protein interactions in TIME and non-TIME drivers. **G**. Proportions of TIME and non-TIME drivers that are part of protein complexes. **H.** Distributions of level 2-9 Reactome pathways^23^ containing TIME and non-TIME drivers. **I.** Proportion of TIME and non-TIME drivers mapping to immune-related pathways as derived from MSigDB^24^ and Reactome^23^. CPI = cancer-promoting inflammation, CYS = cytotoxicity score, FDR = false discovery rate, ICR = immunologic constant of rejection, IS = immune score, TIS = tumour inflammation signature. TCGA abbreviations are listed in **Table S1**. Proportions (**C, G, I**) and distributions (**D, F, H**) were compared using Fisher’s exact test and Wilcoxon rank-sum test, respectively. In (**D**), Benjamini-Hochberg correction for multiple testing was applied.

To characterise the TIME of these samples, we used five gene expression signatures indicative of overall tumour immune infiltration (IS)^17^, cytotoxic anti-tumour infiltration (CYS)^12^, anti-tumour T-helper activity (ICR)^15^, anti-tumour inflammation state (TIS)^18^, and cancer-promoting inflammation (CPI)^19,20^ (**Table S2**). The five gene signatures overlapped only minimally (**Fig. S1 B**), confirming that they captured distinct TIME properties. Since IS and CPI were available only for a subset of samples, we recomputed them for the whole cohort, verifying that our results reproduced those previously published^17,19,20^ (**Fig. S1C, D**).

We grouped samples into low, medium, and high TIME levels based on the corresponding score distribution of each TIME feature in each cancer type. We then calculated the probability of a cancer driver to predict the TIME level of the sample where it was altered using ordinal logistic regression with Lasso regularisation independently for each feature in each cancer type (**Fig. 1A**). Driver-TIME feature pairs with a positive *β* regression coefficient indicated that samples with damaging alterations in that driver were likely to have medium or high levels for that TIME feature. Driver-TIME features pairs with a negative *β* regression coefficients indicated the opposite.

Overall, we identified 477 TIME drivers whose damaging alterations predicted higher (301) or lower (277) TIME levels in 30 cancer types (**Fig. 1A, Table S3**). These predictions included 66 of 102 experimentally validated TIME drivers (p = 4×10^−8^, Fisher’s exact test, **Table S4**), supporting their robustness. Cholangiocarcinoma and kidney chromophobe cancer did not show any significant association, possibly because of the low statistical power due to their small cohorts. Predicted TIME drivers included 164 canonical and 313 candidate drivers, indicating that most well-known cancer drivers can interfere with the immune system (**Fig. 1B**). Unexpectedly since cancer drivers tend to be cancer-specific^16^, TIME drivers were instead recurrently damaged across cancer types (**Fig. 1C**) and samples (**Fig. 1D**, **Table S5**). Moreover, more than 40% of them were also predicted in multiple cancer types (**Table S3**). These included well-known TIME drivers such as *TP53, PTEN, ARID1A*, and *KRAS*, but also *PIK3CA, CDKN2A*, or *TERT* for which no or very little interactions with the TIME have been described. Most TIME drivers (261, 55%) were predictive of at least two features, and 45 of all five of them (**Fig. 1E**). An example was *BRAF*, whose V600E mutation is highly immunogenic^21^, despite BRAF signalling promoting pro-tumour inflammation^22^.

Our results depicted TIME drivers as genes recurrently damaged across cancer samples and types and able to interact plastically with multiple TIME features. This suggested that TIME drivers were likely multifunctional genes involved in several biological processes. To test this hypothesis, we computed the number of interactions of TIME drivers in the protein-protein interaction network. We confirmed that TIME drivers encoded proteins engaging in multiple protein-protein interactions (**Fig. 1F**) and frequently part of protein complexes (**Fig. 1G**). Moreover, TIME drivers mapped to a significantly higher number of biological pathways (**Fig. 1H**) and were involved in a higher number of immune-related functions (**Fig. 1I**) compared to non-TIME drivers.

### TIME tumour suppressors and oncogenes predict opposite TIME states

Given their different mode of action, we sought to analyse the TIME interactions of tumour suppressors and oncogenes separately. Overall, we found that their alterations had an opposite effect on the TIME composition. While tumour suppressors were enriched in TIME drivers predictive of a hot anti-tumour TIME, oncogenes were enriched in TIME drivers predictive of cold pro-tumour TIME (**Fig. 2A, Table S6**). These observations suggested that tumour suppressor alterations preferentially helped tumours to survive in a hot TIME. Oncogene alterations, instead, sustained tumour growth in the presence of a pro-tumour TIME.

**Fig. 2.**
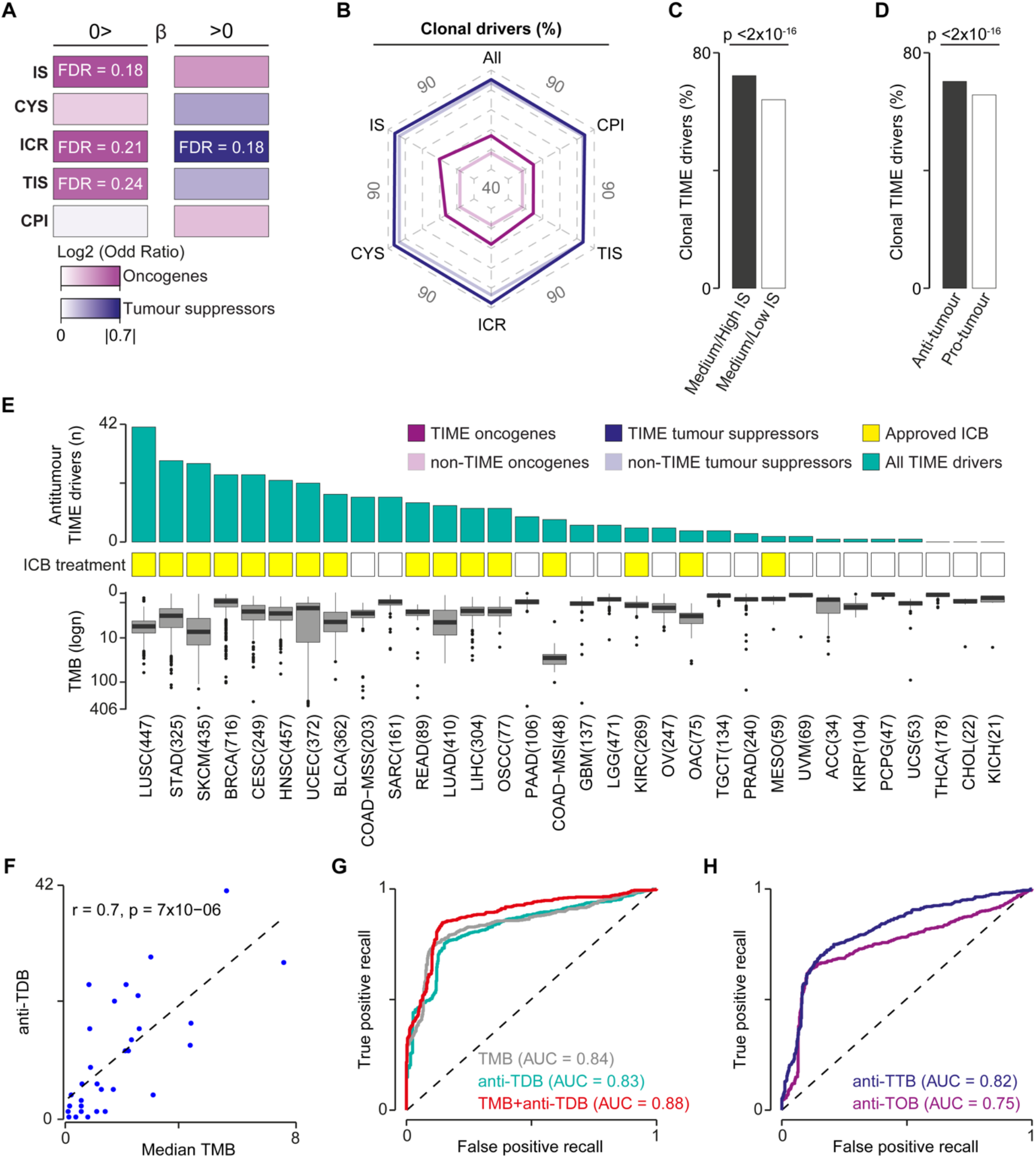
Effects of tumour suppressors and oncogenes on TIME and ICB response. **A.** Enrichment of TIME drivers predictive of medium/low (*β* < 0) or medium/high (*β* > 0) TIME levels in tumours suppressors and oncogenes. **B.** Proportions of TIME and non-TIME tumour suppressors or oncogenes with clonal damaging alterations. **C.** Proportions of clonal TIME drivers predictive of high or low immune infiltration. **D.** Proportions of clonal TIME drivers predictive of an anti-tumour (CYS, TIS, ICR *β >* 0, or CPI *β* < 0) and pro-tumour (CYS, TIS, ICR *β* < 0, or CPI *β* > 0) TIME. **E.** Number of antitumour TIME drivers, approval ICB treatment and TMB across cancer types. The number of samples for each cancer type is shown brackets. **F.** Pearson’s correlations between the median TMB and the number of antitumour TIME drivers in 31 cancer types, excluding COAD-MSI. ROC curves comparing the performance of TMB and anti-TDB (**G**) and TOB or TTB (**H**) in predicting response to ICB. Recall rates and AUCs were calculated across 100 cross-validations. AUC = area under the curve, CPI = cancer-promoting inflammation, CYS = cytotoxicity score, ICB = immune checkpoint blockade, ICR = immunologic constant of rejection, IS = immune score, anti-TDB = antitumour TIME driver burden, TIME = tumour immune microenvironment, TIS = tumour inflammation signature, TMB = tumour mutational burden, TOB = TIME oncogene burden, TTB = TIME tumour suppressor burden. TCGA abbreviations are listed in **Table S1**. Proportions (**A-D**) were compared using Fisher’s exact test. In (**A**), Benjamini-Hochberg correction for multiple testing was applied.

We reasoned that, if alterations in TIME tumour suppressors favoured immune escape, they were likely to occur early in tumour evolution. To test this hypothesis, we computed the proportion of clonal drivers and found that TIME tumour suppressors were enriched in clonal drivers compared to TIME oncogenes and non-TIME tumour suppressors (**Fig. 2B**, **Table S7**). Moreover, the proportion of clonal TIME drivers predictive of high immune infiltration was significantly higher than that of TIME drivers predictive of low immune infiltration (**Fig. 2C**). Similarly, anti-tumour TIME drivers were enriched in clonal alterations compared to pro-tumour TIME drivers (**Fig. 2D**). Therefore, cancers with a hot TIME start to select alterations in tumour suppressors very early in their evolution. Interestingly, also the proportion of TIME oncogenes with clonal alterations was significantly higher than that of non-TIME oncogenes (**Fig. 2B**, **Table S7**). This suggested that, independently on their effect, drivers involved in the tumour-immune interactions are altered earlier than other drivers. Interestingly, 74% of genes driving somatic clonal expansion in non-cancer tissues^25^ were TIME drivers, indicating that their interaction with the immune system may even predate cancer transformation.

A hot TIME is needed for an effective response to immune checkpoint blockade (ICB)^4^. We therefore hypothesised that the number of antitumour TIME drivers in a cancer type (*i. e*. its antitumour TIME driver burden, anti-TDB) could predict its tendency to respond to ICB. To test for this, we considered whether ICB treatment had been approved for that cancer type^26,27^ and used the median tumour mutational burden (TMB) for comparison. Unsurprisingly since both anti-TDB and TMB depend on the overall number of cancer alterations, they were positively correlated (**Fig. 2E, F**). We used Bayesian logistic regression to account for this correlation^28^ and found that TMB and anti-TDB were equally predictive of response to ICB (p = 0. 003). We then compared the predictive power of TMB and anti-TDB alone or in combination using Receiver Operating Characteristic (ROC) curves. We confirmed that TMB and anti-TDB alone were significant predictors of response, but their combination further improved their predictive power (**Fig. 2G**). In line with their antitumour TIME interactions, antitumour TIME tumour suppressor burden (anti-TTB) had higher predictive power than the TIME oncogene burden (anti-TOB, **Fig. 2H**).

### TIME drivers predict the TIME profiles of head and neck cancer subtypes

To gain further insights into driver-TIME interactions, we focused on head and neck squamous cell carcinoma (HNSC), which identifies group of genetically heterogeneous cancers from multiple anatomical sites^29^. The two main subtypes are caused by human papillomavirus (HPV^+^ HNSC) and cigarette smoking (HPV^-^ HNSC), respectively. HPV^+^ tumours show fewer genetic instability, respond better to treatments and have an overall better prognosis^30^. Despite having among the highest leukocyte infiltration^14^, HNSC shows variable response to immunotherapy^26^. This makes it an interesting cancer type to further investigate the dynamic of driver-TIME interactions.

We expanded the TCGA HNSC cohort to include samples from the Clinical Proteomic Tumour Analysis Consortium (CPTAC)^31^, for a total of 562 samples with matched genomic and transcriptomic data (**Fig. 3A**). Of these, 68 were positive to human papillomavirus (HPV^+^). Based on the levels copy number alterations (CNAs)^32^, we further divided the remaining HPV negative (HPV^-^) HNSCs into 351 CNA^high^ and 143 CNA^low^ samples (**Fig. 3A, Fig. S2A**). We confirmed that HPV^+^ HNSC patients have better overall survival (**Fig. 3B**) and, within the HPV^-^ group, high levels of aneuploidy confer worse prognosis^29^ (**Fig. 3C**).

**Fig. 3.**
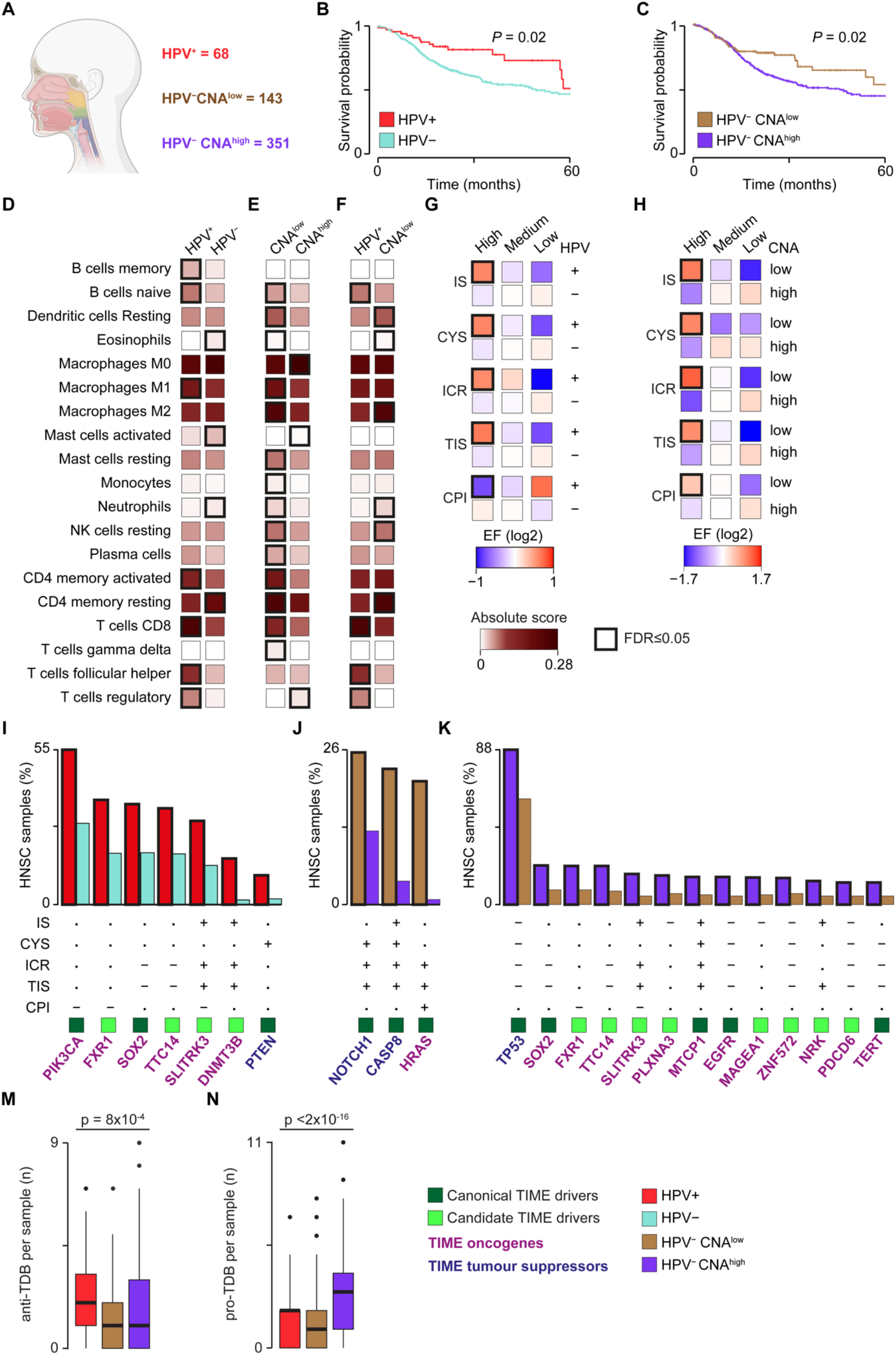
Immune profiles and TIME driver alterations of HNSC. **A.** HNSC extended cohort. HNSCs collected from TCGA and CPTAC were divided in HPV^+^, HPV^-^ CNA^low^ and HPV^-^ CNA^high^ samples based on HPV infection and level of aneuploidy^32^. Kaplan-Meier survival curves between HPV^+^ and HPV^-^ (**B**) or HPV^-^ CNA^low^ and HPV^-^ CNA^high^ (**C**) HNSC patients. Overall survivals were compared using the log-rank test. Comparison of CIBERSORTx absolute score medians between HPV^+^ and HPV^-^ (**D**); HPV^-^ CNA^low^ and HPV^-^ CNA^high^ (**E**); or HPV^+^ and HPV^-^ CNA^low^ (**F**) HNSCs. Only immune cell types enriched in at least one HNSC subtypes are shown. Comparison of enrichment factors (EFs) in the five TIME features between HPV^+^ and HPV^-^ (**G**) or HPV^-^ CNA^low^ and HPV^-^ CNA^high^ (**H**) HNSCs (see **Methods**). TIME drivers more frequently damaged in HPV^+^ HNSCs (**I**), HPV^-^ CNA^low^ (**J**), or HPV^-^ CNA^high^ HNSC samples (**K**). For HPV^-^ CNA^high^ HNSCs only the top 13 TIME drivers are shown (full list in **Table S3**). CPI = cancer-promoting inflammation, CYS = cytotoxicity score, CPTAC = Clinical Proteomic Tumour Analysis Consortium, EF = enrichment factor, FDR = false discovery rate, HPV = human papillomavirus, ICR = immunologic constant of rejection, IS = immune score, TIS = tumour inflammation signature. Proportions were compared using Fisher’s exact test (**D-F, I-K**) or Mantel-Haenszel chi-square test (**G, H**). Distributions (**M, N**) were compared using Kruskal-Wallis test. In (**D-K**), Benjamini-Hochberg correction for multiple testing was applied.

We quantified the immune infiltrates in samples of the three HNSC subtypes from their gene expression profiles and confirmed the hot anti-tumour TIME of HPV^+^ HNSCs (**Fig. 3D**, **Table S8**). Interestingly, we observed an overall higher anti-tumour immunity in HPV^-^ CNA^low^ compared to HPV^-^ CNA^high^ HNSCs (**Fig. 3E**). HPV^-^ CNA^high^ HNSCs were instead enriched in M0 macrophages and regulatory T cells, suggesting that these pro-tumour immune infiltrates could contribute to their worse prognosis. When compared directly, HPV^+^ and CNA^low^ HNSCs showed a different immune infiltration profile, with the former enriched in T cells but depleted in NK cells and neutrophils compared to the latter (**Fig. 3F**). The profiles of the five TIME features across HNSC subtypes confirmed that both HPV^+^ HNSCs and HPV^-^ CNA^low^ HNSCs were rich in anti-tumour and poor in pro-tumour TIME (**Fig. 3G, H**, **Table S9**).

To test whether the TIME features of the three HNSC subtypes could be explained by their TIME driver alteration profile, we compared the frequency of the 53 HNSC TIME drivers across subtypes (**Table S3**). Five of the seven TIME drivers significantly more frequently damaged in HPV^+^ HNSCs were predictive of high anti-tumour or low pro-tumour TIME (**Fig. 3I**). Similarly, all three TIME drivers more frequent in HPV^-^ CNA^low^ HNSCs were predictive of high anti-tumour immunity, while most TIME drivers more frequent in HPV^-^ CNA^high^ were predictive of low anti-tumour immunity (**Fig. 3J, K**). Moreover, HPV^+^ and HPV^-^ CNA^low^ HNSCs showed significantly higher anti-tumour or lower pro-tumour TDB per sample than HPV^-^ CNA^high^ HNSCs (**Fig. 3M, N**).

These results indicated that the distinct immune profiles within HNSCs segregate with the distinct TIME driver alteration profiles across molecular subtypes.

### Functional networks uncover the molecular mechanisms of driver-TIME interactions

To unravel the functional links between driver alterations and TIME features in HNSC, we rebuilt the transcriptional regulatory network of 1,443 transcription factors (TFs) in 562 HNSCs using their expression profiles (**Fig. 4A**). Measuring TF protein activity, we found 1,211 TFs overall active in HNSC and 398 differentially active in the three HNSC subtypes. Of these, 240 showed a significant correlation with TIME features in one of the three HNSC subtypes (TIME TFs). Comparing the protein activity of these TIME TFs in HNSCs with and without damaging alterations in the 53 HNSC TIME drivers, we found 1,386 TIME driver – TIME TF associations. We then combined several types of functional data (**Methods**) and identified seven functional networks linking HNSC TIME drivers and TIME TFs (**Fig. 4A, Tables S10**). Since these networks comprised between 37 and 203 functional nodes (**Table S11**), we extracted the coherent subnetworks connecting TIME drivers to TIME TFs through maximum three nodes.

**Fig. 4.**
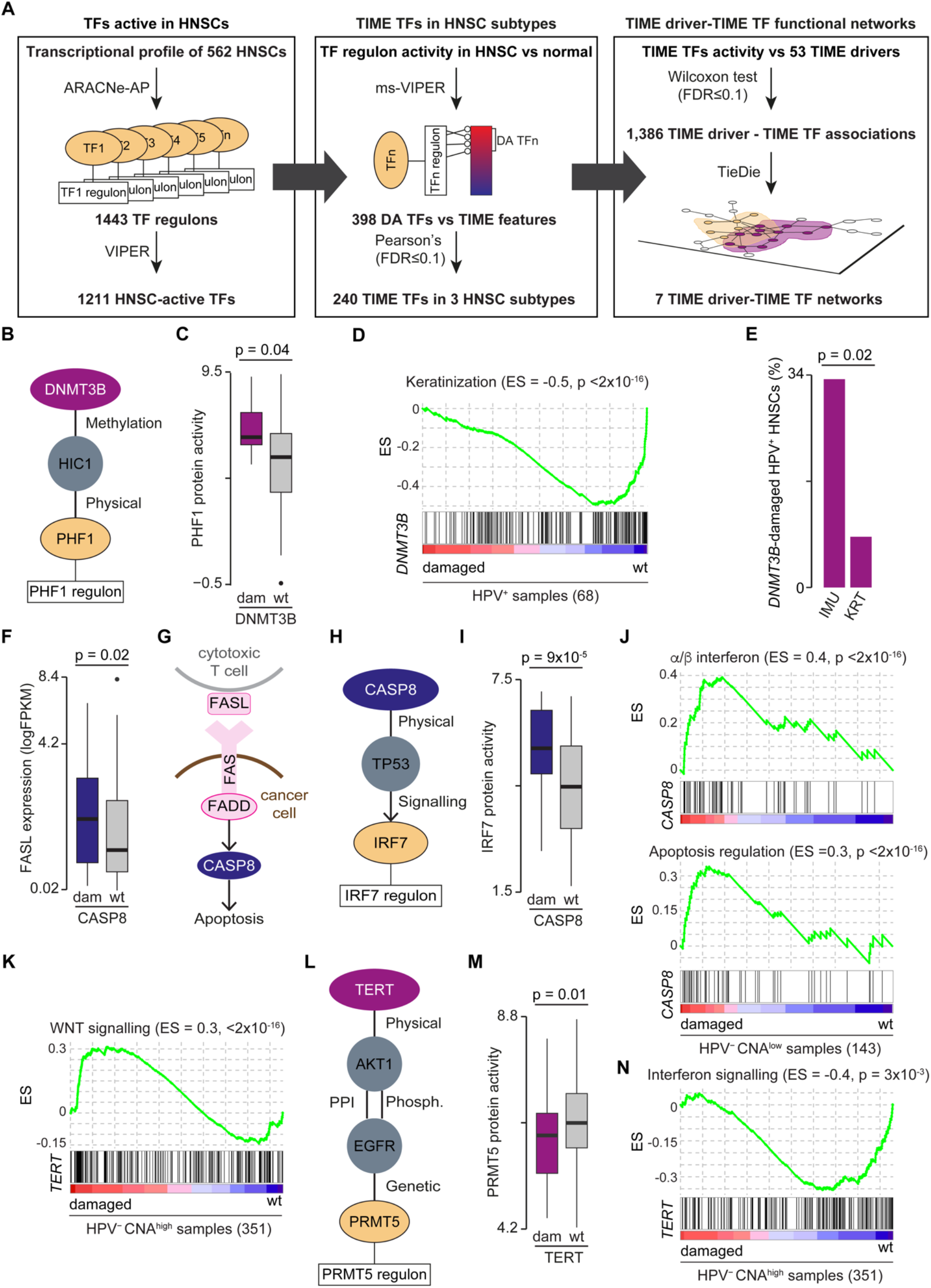
Driver–TIME functional networks in HNSC subtypes. **A.** Reconstruction of HNSC driver-TIME functional networks. HNSC transcriptional regulatory network was used to identify the transcription factors (TFs) differentially active (DA) in the three HNSC subtypes that correlated with TIME features and were associated with TIME drivers. Combining functional data, the significant functional networks linking these drivers to TIME TFs were derived. **B.** DNMT3B functional subnetwork in HPV^+^ HNSCs. **C.** Comparison of PHF1 protein activity between *DNMT3B*-damaged and wild-type (wt) HPV^+^ HNSCs. **D.** Gene set enrichment analysis (GSEA) plot comparing the activation of the keratinization pathway between *DNMT3B*-damaged and wt HPV^+^ HNSCs. **E.** Comparison of *DNMT3B*-damaged samples between immune (IMU) and keratinization (KRT) HPV^+^ HNSCs from^36^. **F.** Comparison of *FASL* gene expression levels between *CASP8*-damaged and wt HPV^-^ CNA^low^ HNSCs. **G.** Schematic of cytotoxic T-cell induced apoptosis of cancer cells through the FAS-FASL cascade. **H.** CASP8 functional subnetwork in HPV^-^ CNA^low^ HNSCs **I.** Comparison of IRF7 protein activity in *CASP8*-damaged and wt HPV^-^ CNA^low^ HNSCs. **J.** GSEA plots comparing the activation of the a/β interferon signalling and apoptosis regulation pathways between *CASP8*-damaged and wt HPV^-^ CNA^low^ HNSCs. **K.** GSEA plot comparing the activation of the WNT signalling pathway between *TERT*-damaged and wt HPV^-^ CNA^high^ HNSCs. **L.** TERT functional subnetwork in HPV^-^ CNA^high^ HNSCs. **M.** Comparison of PRMT5 protein activity between *TERT*-damaged and wt HPV^-^ CNA^high^ HNSCs. **N.** GSEA plot comparing the activation of the interferon signalling pathway between *TERT*-damaged and wt HPV^-^ CNA^high^ HNSCs. CNA = copy number alteration, HNSC = head and neck squamous cell carcinoma, HPV = human papilloma virus, TIME = tumour immune microenvironment. Distributions (**C, F, I, M**) were compared using Wilcoxon rank-sum test. Proportions (**E**) were compared using Fisher’s exact test. GSEAs (**D, J, K, N**) were performed using gene set permutation tests.

These subnetworks enabled investigation of the transcriptional programmes directly linking TIME driver alterations to the TIME features in each HNSC subtype. For example, the TIME oncogene *DNMT3B*, predictive of a hot TIME, was frequently damaged in HPV^+^ HNSCs (**Fig. 3I**). DNMT3B was part of the HPV^+^/TIS subnetwork involving HIC1 and the TIME TF PHF1 (**Fig. 4B, Table S11**). DNMT3B is known to methylate HIC1^33^ inhibiting PHF1 recruitment and activating its transcriptional repression programme^34^. Consistently, we found a higher PHF1 protein activity (**Fig. 4C**) and a downregulation of keratinization (a pathway enriched in PHF1 targets, **Table S12**) in *DNMT3B*-damaged HPV^+^ HNSCs (**Fig. 4D**). Keratinocytes have recently been reported to inhibit T cell proliferation by secreting T cell-modulating cytokines^35^. This could explain how *DNMT3B* amplification could lead to higher immune infiltration. Interestingly, HPV^+^ HNSCs have been divided into two transcriptional subtypes, one (HPV^+^ KRT) characterized by high keratinocyte differentiation and the other (HPV^+^ IMU) with a strong immune response^36^. Using the same dataset^36^, we verified that *DNMT3B* was more frequently damaged in HPV^+^ IMU HNSCs (**Fig. 4E**). This independently supported the hot anti-tumour TIME induced by *DNMT3B* amplification.

Next, we investigated the TIME role of the tumour suppressor *CASP8* whose damaging alterations were predictive of anti-tumour immunity and were enriched in HPV^-^ CNA^low^ HNSCs (**Fig. 3J**). Two lines of evidence supported a role of *CASP8* loss in immune escape in HPV^-^ CNA^low^ HNSC. The first was that *FASL*, a cytotoxic T cell-induced trigger of apoptosis^37^ was upregulated in *CASP8*-damaged HPV^-^ CNA^low^ HNSCs (**Fig. 4F**). Since CASP8 is the downstream target of FASL-initiated apoptotic cascade (**Fig. 4G**), its loss could prevent cancer cells to undergo apoptosis. The second line of evidence came from the HPV^-^ CNA^low^ HNSC subnetworks where CASP8 interacts with the TIME TF IRF7 through TP53 (**Fig. 4H, Table S11**). Since CASP8 loss stabilises TP53^38^, we expected higher IRF7 activity in *CASP8*-damaged HPV^-^ CNA^low^ HNSCs, which was indeed confirmed (**Fig. 4I**). IRF7 targets were enriched in several immune and apoptosis-related pathways (**Table S12**). Accordingly, we found a significant upregulation of both a/β interferon signalling and apoptosis negative control *CASP8*-damaged HPV^-^ CNA^low^ HNSCs (**Fig. 4J**). This further confirmed that apoptosis reduction was a CASP8-induced immune escape mechanism in HPV^-^ CNA^low^ HNSCs with a hot TIME.

Lastly, we analysed the functional network of the TIME oncogene *TERT*, predictive of a cold TIME and whose GoF alterations were enriched in HPV^-^ CNA^high^ HNSCs (**Fig. 3K**). *TERT* is a member of the WNT-β-catenin pathway and its activation led to WNT upregulation in *TERT*-damaged HPV^-^ CNA^high^ HNSCs (**Fig. 4K**). WNT activation has been linked to immune exclusion^8^, which could explain the role of TERT in inducing a cold TIME. Moreover, TERT was in the same HPV^-^ CNA^high^ subnetwork of the TIME TF PRMT5 (**Fig. 4L, Table S11**). TERT activation is known to induce AKT1-mediated EGFR phosphorylation that, in turn, downregulates PRMT5 through a negative genetic interaction^39^. We confirmed a lower PRMT5 protein activity (**Fig. 4M**) and the downregulation of interferon signalling (**Fig. 4N**), one of the pathways enriched in PRMT5 targets (**Table S12**) in *TERT*-damaged HPV^-^ CNA^high^ HNSCs. A lower interferon activity could reduce the production of T cell chemo-attractants resulting in a cold TIME.

## DISCUSSION

In this study, we predicted the functional interactions between the genetic drivers of 6,921 cancers and their immune microenvironment. Despite the analysis being conducted separately in 30 cancer types, the predicted TIME drivers shared key properties, including high multifunctionality, plasticity in their interaction with the TIME, and recurrent damaging alterations across cancer types and samples. These properties support a multifaceted role of TIME drivers in promoting tumour evolution through both cancer-intrinsic and cancer-extrinsic mechanisms and suggest that they can interfere with multiple TIME features likely in a tissue-specific manner.

We found an enrichment of TIME tumour suppressors in early mutated drivers and their alterations are predictive of a hot anti-tumour TIME. These observations strongly suggesting that they are involved in immune evasion. This agrees with the recently reported preferential loss of tumour suppressors in mouse models with a functional adaptive immunity^10^ and supports their emergent role as the guardians of immune integrity^40^. Differently from tumour suppressors, TIME oncogenes were preferentially damaged in samples with a cold TIME, in line with the documented role of *MYC, HRAS* and *BRAF* oncogenes in inducing inflammatory chemokines and cytokines^41^. The opposite effect of tumour suppressors and oncogenes on the TIME could reflect their broad functional differences, with the former mostly involved in controlling cell cycle, DNA repair and apoptosis and the latter enriched in signalling genes^42^. In all cases, TIME alterations occur earlier than those of other drivers, suggesting that the interactions between the mutated epithelium and the immune cell compartment is likely to start well before the epithelial cells become fully transormed^25^.

The burden of antitumour TIME drivers, particularly tumour suppressors, can predict whether a cancer type is responsive to ICB similarly to TMB and their combination improves the overall predictive power. The identification of patients who are most likely to benefit from ICB treatment is an open clinical question since response to ICB primarily assessed in the clinic and translational biomarkers are still very few^43^. For example, although ICB treatment is now standard of care in recurrent HNSC^44^, the majority of patients will not respond^45^ exposing them to unnecessary toxic effects and worse survival. In clinical practice, the combined positive score based on the number of PD-L1 positive cells over all tumour cells is used to determine eligibility to ICB treatment^46^. Still, only 30% of HNSC patients will respond, some of which with dramatic and prolonged results but currently there are no reliable biomarkers^26^. We showed that HPV^+^ but also HPV^-^ CNA^low^ HNSCs tend to have a hot anti-tumour TIME while HPV^-^ CNA^high^ HNSCs are usually deprived of immune infiltration. This would suggest a prioritisation of ICB treatment only in patients with HPV^+^ and HPV^-^ CNA^low^ HNSC subtypes.

Unlike TMB that has a sample-specific value, anti-TDB is a feature of the cancer type and cannot predict response to ICB of the individual patient. To overcome this limitation and unravel the molecular mechanisms of the driver-TIME interactions, we rebuilt the transcription regulatory networks linking directly driver alterations to TIME states. We identified TIME-driver functional networks for 33 HNSC TIME drivers, indicating how alterations in these genes interfere with the TIME. For example, *DNMT3B*-damaged HPV^+^ HNSCs significantly overlap with the recently identified HPV^+^ IMU subtype, which shows better prognosis and high immune infiltration^36^. Our data show that this is likely achieved through a reduction of keratinocyte differentiation induced by *DNMT3B* amplification. Therefore, patients with *DNMT3B*-damaged HPV^+^ HNSCs are good candidates for a successful ICB treatment. Similarly, CASP8-induced immune escape through apoptotic inhibition is another mechanism evolved by HPV^-^ CNA^low^ HNSCs to survive a high anti-tumour infiltration. Our prediction is that also this subgroup of patients would benefit from ICB treatment. On the contrary, TERT activation modulates a cold TIME, suggesting that HPV^-^ CNA^high^ patients with this alteration would not benefit from immunotherapy.

Besides these examples, our analysis provides a comprehensive set of driver-TIME interactions and mechanistic insights into their crosstalk that could be further explored in experimental and clinical settings.

## METHODS

### TCGA data

A dataset of 7730 TCGA samples with quality-controlled mutation (SNVs and indels), copy number and gene expression data in 32 solid cancer types was assembled from the Genomic Data Commons portal (GDC, https://portal.gdc.cancer.gov/). Oesophageal cancer was divided into squamous cell carcinoma (OSCC) and adenocarcinoma (OAC) using TCGA annotation. Colon adenocarcinoma was split into COAD-MSS and COAD-MSI according to the level of microsatellite instability (MSI)^47^.

All SNVs and indels were annotated with ANNOVAR^48^ (April 2018) and dbNSFP^49^ v3. 0 and only those identified as damaging were retained. These included truncating mutations (stopgain, stoploss, frameshift), hotspot mutations, missense mutations and splicing mutations predicted as damaging as described in^16^.

Copy Number Alteration (CNA) segments, sample ploidy and sample purity were obtained from TCGA SNP arrays using ASCAT^50^ v. 2. 5. 2. Segments were intersected with the exonic coordinates of 19,641 unique human genes in hg19 and CNA genes were identified as those with ≥25% of transcribed length covered by a CNA segment. RNA-Seq data were used to filter out CNAs with no effect on gene expression. Damaging gene gains were defined as CN>2 times sample ploidy and significant increase in gene expression as compared to baseline expression using a Wilcoxon rank-sum test and accounting for multiple testing using Benjamini-Hochberg correction. Only gene gains with false discovery rate (FDR) <0.05 were retained. Homozygous gene losses were defined as CN = 0 and FPKM expression values <1 over sample purity. Heterozygous gene losses were defined as CN = 1 or CN = 0 but FPKM expression values >1 over sample purity. This resulted in 2,163,756 redundant genes damaged in 7,730 TCGA samples. Of these, 511,048 genes acquired LoF alterations (homozygous deletion, truncating, missense damaging, splicing mutations, or double hits), while 1,652,708 genes were considered to acquire GoF alterations (hotspot mutation or damaging gene gain).

### Driver annotation

The cancer drivers for each of 32 cancer types were retrieved from the Network of Cancer Genes (NCG, http://www.network-cancer-genes.org), which collects preferentially mutational drivers^16^. To add drivers altered through CNAs, focal amplifications and deletions in each cancer type were gathered from the FireBrowse portal (http://firebrowse.org)^51^. Amplification and deletion segments were intersected with 256 canonical and 1,405 candidate oncogenes and 254 canonical and 1,318 candidate tumour suppressors^16^, respectively. Drivers for at least 25% of their transcript within a CNA event were considered CNA drivers. Only tumour suppressors with LoF alterations (homozygous deletion, truncating, missense damaging, splicing mutations, or double hits) and oncogenes with GoF alterations (hotspot mutation or damaging gene gain) were retained. Both LoF and GoF alterations were considered for drivers with unclassified roles. Finally, only drivers damaged in ≥2% or five samples were retained. In total 1,231 (254 canonical and 977 candidate drivers, **Fig. 1 and S1A**) damaged in 6,921 samples were used for the identification of TIME drivers.

The clonality of 27,763 damaging mutations affecting 1231 drivers was measured using the cancer cell fraction (CCF) as described in^52^. Briefly, for each damaging mutation, the probability to have a CCF from 0. 01 to 1 incremented by 0. 01 was calculated given the observed variant allele frequency (VAF), copy number status of the mutation, sample purity and normal copy number. The CCF with the highest probability was selected with the associated 95% confidence interval (CI). A damaging mutation was considered clonal if 95% CI of the CCF overlapped with 1, otherwise it was considered subclonal. A driver was considered clonal when it had at least one clonal damaging mutation.

The clonality of 38,846 damaging CNAs affecting 1231 drivers was assessed with ABSOLUTE^53^, using mutation VAFs and SNP6 array segmentation values obtained from GDC. ABSOLUTE was run with default parameters and the cancer type-specific karyotype models for 6,900 TCGA samples. Even in this case the returned CCF with the highest probability was selected with the associated 95% CI. A CNA driver was considered clonal if a 95% CI of its CCF overlapped with 1, otherwise it was considered subclonal.

### TIME features

To assess the cytotoxic anti-tumour infiltration score (CYS), 6,445 samples were grouped into six clusters ordered from the lowest (CYS1) to the highest (CYS6) CYS levels, as defined in the original publication^12^. The six CYS levels were grouped into low (CYS1, 2), medium (CYS 3, 4) and high (CYS 5, 6) groups for consistency with the other features. To assess the immunologic constant of rejection (ICR), 6528 samples were grouped into low, medium or high ICR levels based on the expression distribution of 20 genes encoding IFN-simulated, regulatory, and effector immune molecules^15^. To assess the tumour inflammation signature (TIS), 6266 samples were grouped into low, medium or high TIS levels based on the expression values of 18 genes measuring adaptive immune response^18^. Overall immune infiltration (immune score, IS) and cancer-promoting inflammation (CPI) values were calculated for 6921 and 6728 samples, respectively, based on the expression of 141 genes using ESTIMATE^17^ and as the mean of the Log2-transformed expression^20^ of ten genes encoding known mediators of cancer-promoting inflammation^19^, respectively. For IS, FPKM gene expression data from GDC were used to assess gene expression levels. For CPI, RSEM gene expression data from cBioPortal^54,55^ were used. Cancer samples were grouped into discrete categorical levels starting the lowest (low) to the highest quartile (high) and assigning all remaining samples to the group with medium levels. The sample grouping was performed for each cancer type separately.

### Lasso-regularised ordinal regression

Lasso-regularised ordinal regression was used to estimate the probability of each damaged driver to predict the TIME ordinal level of each sample using the glmnetcr function from the glmnetcr R package^56^ v. 1. 0. 6. The input data consisted of a binary matrix whose rows corresponded to samples (observations) and columns (variable) to the driver alteration status. TIME levels were encoded as ordered factor vectors with a size equal to the number of samples. Regression analysis was run for each TIME feature in each cancer type, considering canonical and candidate drivers separately and only samples with ≥1 damaged driver, for a total of 320 glmnetcr calls to fit 320 regression models. All analyses were run without variable standardisation and all other parameters set to default values. In each regression, multiple steps of the model with the different values of lambda were ran and models with the minimum Akaike information criterion (AIC) were used to extract non-zero *β* coefficients^56^.

### Protein-protein interaction and functional analysis

The number of non-redundant protein-protein interactions for TIME and non-TIME drivers were computed from BioGRID^57^ v. 3. 5. 185, IntAct^58^ v. 4. 2. 14, DIP^59^ (February 2018), HPRD^60^ v. 9 and Bioplex^61^ v. 3. 0 as described in^16^ and compared using a Wilcoxon rank-sum test. The proportion of proteins encoded by TIME and non-TIME cancer drivers that engage in complexes were derived from CORUM^62^ v. 3. 0, HPRD^60^ v. 9 and Reactome^23^ v. 72 as described in^16^ and compared using Fisher’s exact test.

Reactome^23^ v. 72 level 2-9 pathways were used to calculate the numbers of pathways each of 821 drivers present in Reactome mapped to. These were compared between 335 TIME and 486 non-TIME cancer using a Wilcoxon rank-sum test. A list of 2,519 immune-related genes was derived combining genes mapping to the immune system level 1 pathway of Reactome^23^ v. 72 and the immune-related pathways in MSigDB^24^. The proportions of immune-related TIME and non-TIME drivers were compared using a Fisher’s exact test.

### HNSC extended cohort

A dataset of 109 HNSC samples from the Clinical Proteomic Tumour Analysis Consortium (CPTAC) with quality-controlled mutation (SNVs and indels), copy number and gene expression data was downloaded from GDC. Damaging mutations were identified as described above. CNAs were derived using AscatNGS^63^. Sample ploidy was calculated as the average copy number of all segments weighted by segment length^63^. Sample purity was measured from gene expression data using ESTIMATE^17^. Gene gains, heterozygous and homozygous gene losses were defined as described above. In total, 26,450 redundant genes were damaged with 7,891 LoF and 18,559 GoF alterations. The CPTAC samples were added to the TCGA cohort for a total of 562 HNSCs.

HPV^-^ HNSCs were divided into CNA^high^ and CNA^low^ subtypes as described in^32^ using a cohort of 1,495 squamous cell carcinomas that included 1,386 TCGA samples and 109 CPTAC HNSCs. CNA GISTIC2 loci were obtained from^32^ for the TCGA cohort and from LinkedOmics^64^ for the CPTAC HNSCs. Loci were classified as copy number neutral, low and high CNAs and used for hierarchical clustering using Euclidean distance. Two clusters, one with 143 HPV^-^ CNA^low^ HNSCs and the other with 351 HPV^-^ CNA^high^ HNSCs were identified. TCGA classification overlapped with that in the original publication^32^ for 94% of samples (**Figure S2A**). Survival analysis was performed for 557 patients with available clinical data using the survminer R package v. 0. 4. 9 and compared between HNSC subtypes using the log-rank test.

CIBERSORTx^65^ was run on the FPKM-normalised RNA-seq data of 562 HNSCs using the LM22 signature to estimate the absolute abundance level of 22 immune populations. Absolute abundance scores were compared between HNSC subtypes using a Wilcoxon rank-sum test and corrected with Benjamini-Hochberg correction.

Since only FPKM gene expression data were available for all 562 HNSCs, the five TIME features were recalculated using FPKM instead of RSEM values, verifying that the two measures correlated positively (**Figure S2B-F**). For CYS and ICR, clustering was done as described in the original publications^12,15^. For TIS the same clustering strategy as for the TCGA cohort was applied. For IS and CPI, the score was calculated as described above.

For each TIME feature, the enrichment factor (*EF*) of a HNSC subtype *i* in a TIME level *j* (*EF_ij_*) was calculated as:

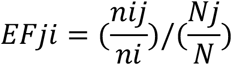

Where *n_ij_* is the number of HNSCs in subtype *i* with TIME level *j; n_i_* is the total number of HNSCs in subtype *i*; *N_j_* is the number of all HNSCs with TIME level *j;* and N is the total number of HNSCs.

### TIME drivers – TIME TFs functional network

A list of 1,471 genes annotated with the GO:0006355 (regulation of DNA-templated transcription) Gene Ontology (release 2022-05)^66,67^ term were considered *bona fide* transcription factors (TFs) and used as input for ARACNE-AP^68^ together with the gene expression profiles of 562 HNSCs with default parameters (**Fig. 4A**). The resulting HNSC transcriptional regulatory network composed of 1,443 TFs, 18,067 targets and 202,512 interactions was used to infer the sample-level TF protein activity using VIPER^69^, resulting in 1,211 HNSC-active TFs.

The expression levels of downstream targets of each active TFs were compared between HNSC subtypes and 100 adjacent normal tissues using ms-VIPER^69^. Overall, 271, 113 and 212 differentially active (DA) TFs (p-value <0. 05) were found in HPV^+^, HPV^-^ CNA^low^ and HPV^-^ CNA^high^ HNSCs, respectively, for a total of 398 unique DA TFs. Pearson’s correlation was calculated between the protein activity of each DA TF and each TIME feature to retrieve DA TFs significantly correlated with TIME in each HNSC subtype (TIME TFs, FDR <0. 1). Overall, 51, 103 and 159 TIME TFs were found in HPV^+^, HPV^-^ CNA^low^ and HPV^-^ CNA^high^ HNSCs, respectively, for a total of 240 unique DA TFs.

TIME TFs were tested for statistical association with 53 HNSC TIME drivers, comparing their protein activity in HNSCs with and without TIME driver alterations using Wilcoxon rank sum test. Overall, 131, 373 and 882 TIME TF-TIME driver significant associations (FDR <0. 1) were found in HPV^+^, HPV^-^ CNA^low^ and HPV^-^ CNA^high^ HNSC, respectively, for a total of 1386 associations. TieDIE^70^ was applied to find functional interactions between significantly associated TIME TFs and HNSC TIME drivers. The prior knowledge network (PKN) for TieDIE was assembled from 542,397 protein-protein^16^, 12,730 phosphorylation^71^, 15,104 genetic^57^ and 34,877 signalling interactions^72^ across 18,053 human genes. Fourteen TIME drivers–TIME TFs functional networks were rebuilt in each HNSC subtype and TIME feature, seven of which had an influence score significantly higher (p-value <0. 08) than random networks with the same degree distribution (**Table S10**). Starting from these networks, coherent subnetworks were defined as those with maximum three nodes between the TIME driver and the TIME TF and a positive coherency score^70^ (**Table S11**).

TIME TF targets were functionally annotated using pathway enrichment analysis as described in^73^ (**Table S12**). GSEA^74^ v4. 3. 2 was used with a gene set permutation test of 1,000 iterations.

## DATA AND CODE AVAILABILITY

The data generated in this study are available within the article and its supplementary data files.

## AUTHOR CONTRIBUTIONS

HM: conceptualised the functional network analysis, conducted the computational analyses and wrote the manuscript. RK: conducted the computational analyses. JPJ: analysed the HNSC data. FDC: conceptualised the study, analysed the data and wrote the manuscript.

## ACKNOWLEDGMENTS

This work was supported by Cancer Research UK [C43634/A25487 to F. D. C.] and [EDDPJT-Nov21\100010 to F. D. C], the Cancer Research UK City of London Centre [C7893/A26233 to F. D. C], innovation programme under the Marie Skłodowska-Curie grant agreement No CONTRA-766030, a donation from the Charles Wilson & Rowena Olegari Foundation and Guy’s Cancer Charity, and the Francis Crick Institute, which receives its core funding from Cancer Research UK (FC001002), the UK Medical Research Council (FC001002), and the Wellcome Trust (FC001002). For the purpose of Open Access, the author has applied a CC BY public copyright licence to any Author Accepted Manuscript version arising from this submission.

## REFERENCES

1. McGranahan, N. & Swanton, C. Clonal Heterogeneity and Tumor Evolution: Past, Present, and the Future. Cell 168, 613–628 (2017).

2. Wellenstein, M.D. & de Visser, K.E. Cancer-Cell-Intrinsic Mechanisms Shaping the Tumor Immune Landscape. Immunity 48, 399–416 (2018).

3. Polyak, K., Haviv, I. & Campbell, I.G. Co-evolution of tumor cells and their microenvironment. Trends Genet 25, 30–8 (2009).

4. O’Donnell, J.S., Teng, M.W.L. & Smyth, M.J. Cancer immunoediting and resistance to T cell-based immunotherapy. Nature reviews Clinical oncology 16, 151–167 (2019).

5. Binnewies, M. et al. Understanding the tumor immune microenvironment (TIME) for effective therapy. Nature medicine 24, 541–550 (2018).

6. Ghosh, M. et al. Mutant p53 suppresses innate immune signaling to promote tumorigenesis. Cancer Cell 39, 494–508.e5 (2021).

7. Liao, W. et al. KRAS-IRF2 Axis Drives Immune Suppression and Immune Therapy Resistance in Colorectal Cancer. Cancer Cell 35, 559–572.e7 (2019).

8. Spranger, S., Bao, R. & Gajewski, T.F. Melanoma-intrinsic β-catenin signalling prevents anti-tumour immunity. Nature 523, 231–235 (2015).

9. Kumagai, S. et al. An Oncogenic Alteration Creates a Microenvironment that Promotes Tumor Progression by Conferring a Metabolic Advantage to Regulatory T Cells. Immunity 53, 187–203.e8 (2020).

10. Martin, T.D. et al. The adaptive immune system is a major driver of selection for tumor suppressor gene inactivation. Science 373, 1327–1335 (2021).

11. Lawson, K.A. et al. Functional genomic landscape of cancer-intrinsic evasion of killing by T cells. Nature 586, 120–126 (2020).

12. Tamborero, D. et al. A Pan-cancer Landscape of Interactions between Solid Tumors and Infiltrating Immune Cell Populations. Clin Cancer Res 24, 3717–3728 (2018).

13. Rooney, M.S., Shukla, S.A., Wu, C.J., Getz, G. & Hacohen, N. Molecular and genetic properties of tumors associated with local immune cytolytic activity. Cell 160, 48–61 (2015).

14. Thorsson, V. et al. The Immune Landscape of Cancer. Immunity 48, 812–830.e14 (2018).

15. Roelands, J. et al. Oncogenic states dictate the prognostic and predictive connotations of intratumoral immune response. J Immunother Cancer 8(2020).

16. Dressler, L. et al. Comparative assessment of genes driving cancer and somatic evolution in non-cancer tissues: an update of the Network of Cancer Genes (NCG) resource. Genome Biol 23, 35 (2022).

17. Yoshihara, K. et al. Inferring tumour purity and stromal and immune cell admixture from expression data. Nat Commun 4, 2612 (2013).

18. Danaher, P. et al. Pan-cancer adaptive immune resistance as defined by the Tumor Inflammation Signature (TIS): results from The Cancer Genome Atlas (TCGA). J Immunother Cancer 6, 63 (2018).

19. Zelenay, S. et al. Cyclooxygenase-Dependent Tumor Growth through Evasion of Immunity. Cell 162, 1257–70 (2015).

20. Bonavita, E. et al. Antagonistic Inflammatory Phenotypes Dictate Tumor Fate and Response to Immune Checkpoint Blockade. Immunity 53, 1215–1229.e8 (2020).

21. Ilieva, K.M. et al. Effects of BRAF mutations and BRAF inhibition on immune responses to melanoma. Mol Cancer Ther 13, 2769–83 (2014).

22. Conciatori, F. et al. BRAF status modulates Interelukin-8 expression through a CHOP-dependent mechanism in colorectal cancer. Commun Biol 3, 546 (2020).

23. Jassal, B. et al. The reactome pathway knowledgebase. Nucleic Acids Research 48, D498–D503 (2020).

24. Liberzon, A. et al. The Molecular Signatures Database Hallmark Gene Set Collection. Cell Systems 1, 417–425 (2015).

25. Acha-Sagredo, A., Ganguli, P. & Ciccarelli, F.D. Somatic variation in normal tissues: friend or foe of cancer early detection? Annals of Oncology 33, 1239–1249 (2022).

26. Morad, G., Helmink, B.A., Sharma, P. & Wargo, J.A. Hallmarks of response, resistance, and toxicity to immune checkpoint blockade. Cell 184, 5309–5337 (2021).

27. Sharma, P. et al. The Next Decade of Immune Checkpoint Therapy. Cancer Discovery 11, 838–857 (2021).

28. Gelman, A., Jakulin, A., Pittau, M.G. & Su, Y.-S. A weakly informative default prior distribution for logistic and other regression models. The Annals of Applied Statistics 2, 1360–1383, 24 (2008).

29. Leemans, C.R., Snijders, P.J.F. & Brakenhoff, R.H. The molecular landscape of head and neck cancer. Nat Rev Cancer 18, 269–282 (2018).

30. Du, E. et al. Long-term Survival in Head and Neck Cancer: Impact of Site, Stage, Smoking, and Human Papillomavirus Status. Laryngoscope 129, 2506–2513 (2019).

31. Huang, C. et al. Proteogenomic insights into the biology and treatment of HPV-negative head and neck squamous cell carcinoma. Cancer Cell 39, 361–379.e16 (2021).

32. Campbell, J.D. et al. Genomic, Pathway Network, and Immunologic Features Distinguishing Squamous Carcinomas. Cell Rep 23, 194–212.e6 (2018).

33. Zeng, S. et al. HIC1 epigenetically represses CIITA transcription in B lymphocytes. Biochim Biophys Acta 1859, 1481–1489 (2016).

34. Boulay, G. et al. Hypermethylated in cancer 1 (HIC1) recruits polycomb repressive complex 2 (PRC2) to a subset of its target genes through interaction with human polycomb-like (hPCL) proteins. J Biol Chem 287, 10509–10524 (2012).

35. Seiringer, P. et al. Keratinocytes Regulate the Threshold of Inflammation by Inhibiting T Cell Effector Functions. Cells 10(2021).

36. Zhang, Y. et al. Subtypes of HPV-Positive Head and Neck Cancers Are Associated with HPV Characteristics, Copy Number Alterations, PIK3CA Mutation, and Pathway Signatures. Clin Cancer Res 22, 4735–45 (2016).

37. Andersen, M.H., Schrama, D., Thor Straten, P. & Becker, J.C. Cytotoxic T cells. J Invest Dermatol 126, 32–41 (2006).

38. Müller, I. et al. Cancer Cells Employ Nuclear Caspase-8 to Overcome the p53-Dependent G2/M Checkpoint through Cleavage of USP28. Mol Cell 77, 970–984.e7 (2020).

39. Vichas, A. et al. Integrative oncogene-dependency mapping identifies RIT1 vulnerabilities and synergies in lung cancer. Nat Commun 12, 4789 (2021).

40. Muñoz-Fontela, C., Mandinova, A., Aaronson, S.A. & Lee, S.W. Emerging roles of p53 and other tumour-suppressor genes in immune regulation. Nat Rev Immunol 16, 741–750 (2016).

41. Mantovani, A., Allavena, P., Sica, A. & Balkwill, F. Cancer-related inflammation. Nature 454, 436–444 (2008).

42. Repana, D. et al. The Network of Cancer Genes (NCG): a comprehensive catalogue of known and candidate cancer genes from cancer sequencing screens. Genome Biol 20, 1 (2019).

43. Ganesan, S. & Mehnert, J. Biomarkers for Response to Immune Checkpoint Blockade. Annual Review of Cancer Biology 4, 331–351 (2020).

44. Cohen, E.E.W. et al. The Society for Immunotherapy of Cancer consensus statement on immunotherapy for the treatment of squamous cell carcinoma of the head and neck (HNSCC). Journal for ImmunoTherapy of Cancer 7, 184 (2019).

45. Napolitano, M., Schipilliti, F.M., Trudu, L. & Bertolini, F. Immunotherapy in head and neck cancer: The great challenge of patient selection. Crit Rev Oncol Hematol 144, 102829 (2019).

46. Crosta, S. et al. PD-L1 Testing and Squamous Cell Carcinoma of the Head and Neck: A Multicenter Study on the Diagnostic Reproducibility of Different Protocols. Cancers (Basel) 13 (2021).

47. Liu, Y. et al. Comparative Molecular Analysis of Gastrointestinal Adenocarcinomas. Cancer Cell 33, 721–735.e8 (2018).

48. Wang, K., Li, M. & Hakonarson, H. ANNOVAR: functional annotation of genetic variants from high-throughput sequencing data. Nucleic Acids Res 38, e164 (2010).

49. Liu, X., Wu, C., Li, C. & Boerwinkle, E. dbNSFP v3.0: A One-Stop Database of Functional Predictions and Annotations for Human Nonsynonymous and Splice-Site SNVs. Hum Mutat 37, 235–41 (2016).

50. Van Loo, P. et al. Allele-specific copy number analysis of tumors. Proc Natl Acad Sci U S A 107, 16910–5 (2010).

51. Mermel, C.H. et al. GISTIC2.0 facilitates sensitive and confident localization of the targets of focal somatic copy-number alteration in human cancers. Genome Biol 12, R41 (2011).

52. Goh, G., McGranahan, N. & Wilson, G.A. Computational Methods for Analysis of Tumor Clonality and Evolutionary History. Methods Mol Biol 1878, 217–226 (2019).

53. Carter, S.L. et al. Absolute quantification of somatic DNA alterations in human cancer. Nat Biotechnol 30, 413–21 (2012).

54. Gao, J. et al. Integrative Analysis of Complex Cancer Genomics and Clinical Profiles Using the cBioPortal. Science Signaling 6, pl1–pl1 (2013).

55. Cerami, E. et al. The cBio Cancer Genomics Portal: An Open Platform for Exploring Multidimensional Cancer Genomics Data. Cancer Discovery 2, 401–404 (2012).

56. Archer, K.J. & Williams, A.A.A. L1 penalized continuation ratio models for ordinal response prediction using high-dimensional datasets. Statistics in medicine 31, 1464–1474 (2012).

57. Oughtred, R. et al. The BioGRID interaction database: 2019 update. Nucleic Acids Research 47, D529–D541 (2019).

58. Orchard, S. et al. The MIntAct project—IntAct as a common curation platform for 11 molecular interaction databases. Nucleic Acids Research 42, D358–D363 (2014).

59. Salwinski, L. et al. The Database of Interacting Proteins: 2004 update. Nucleic Acids Research 32, D449–D451 (2004).

60. Keshava Prasad, T.S. et al. Human Protein Reference Database—2009 update. Nucleic Acids Research 37, D767–D772 (2009).

61. Huttlin, E.L. et al. Dual proteome-scale networks reveal cell-specific remodeling of the human interactome. Cell 184, 3022–3040.e28 (2021).

62. Giurgiu, M. et al. CORUM: the comprehensive resource of mammalian protein complexes—2019. Nucleic Acids Research 47, D559–D563 (2019).

63. Raine, K.M. et al. ascatNgs: Identifying Somatically Acquired Copy-Number Alterations from Whole-Genome Sequencing Data. Curr Protoc Bioinformatics 56, 15.9.1–15.9.17 (2016).

64. Vasaikar, S.V., Straub, P., Wang, J. & Zhang, B. LinkedOmics: analyzing multi-omics data within and across 32 cancer types. Nucleic Acids Res 46, D956–d963 (2018).

65. Newman, A.M. et al. Determining cell type abundance and expression from bulk tissues with digital cytometry. Nature Biotechnology 37, 773–782 (2019).

66. Ashburner, M. et al. Gene ontology: tool for the unification of biology. The Gene Ontology Consortium. Nat Genet 25, 25–9 (2000).

67. The Gene Ontology resource: enriching a GOld mine. Nucleic Acids Res 49, D325–d334 (2021).

68. Lachmann, A., Giorgi, F.M., Lopez, G. & Califano, A. ARACNe-AP: gene network reverse engineering through adaptive partitioning inference of mutual information. Bioinformatics 32, 2233–5 (2016).

69. Alvarez, M.J. et al. Functional characterization of somatic mutations in cancer using network-based inference of protein activity. Nat Genet 48, 838–47 (2016).

70. Paull, E.O. et al. Discovering causal pathways linking genomic events to transcriptional states using Tied Diffusion Through Interacting Events (TieDIE). Bioinformatics 29, 2757–64 (2013).

71. Hornbeck, P.V. et al. PhosphoSitePlus, 2014: mutations, PTMs and recalibrations. Nucleic Acids Res 43, D512–20 (2015).

72. Csabai, L. et al. SignaLink3: a multi-layered resource to uncover tissue-specific signaling networks. Nucleic Acids Res 50, D701–d709 (2022).

73. Mourikis, T.P. et al. Patient-specific cancer genes contribute to recurrently perturbed pathways and establish therapeutic vulnerabilities in esophageal adenocarcinoma. Nat Commun 10, 3101 (2019).

74. Subramanian, A. et al. Gene set enrichment analysis: a knowledge-based approach for interpreting genome-wide expression profiles. Proc Natl Acad Sci U S A 102, 15545–50 (2005).

